# A Beefy-R culture medium: replacing albumin with rapeseed protein isolates

**DOI:** 10.1101/2022.09.02.506409

**Authors:** Andrew J. Stout, Miriam L. Rittenberg, Michelle Shub, Michael K. Saad, Addison B. Mirliani, David L. Kaplan

## Abstract

The development of cost-effective serum-free media is essential for the economic viability of cultured meat. A key challenge facing this goal is high-cost recombinant albumin that is necessary in some available serum-free media formulations. As such, there is substantial interest in finding albumin alternatives which are low-cost, effective, scalable, sustainable, and suitable for food applications. Recently, a serum-free medium termed Beefy-9 was developed for bovine satellite cells (BSCs), which relied on recombinant albumin as a key component to replace fetal bovine serum. Here we alter Beefy-9 by replacing albumin with rapeseed protein isolate, a bulk-protein solution obtained from agricultural waste-streams through simple isoelectric protein precipitation. This new medium, termed Beefy-R, improves BSC growth compared with Beefy-9 while maintaining cell phenotype and myogenicity. These results offer an effective, low-cost, and sustainable alternative to albumin for serum-free culture of muscle stem cells, thereby addressing a key hurdle facing cultured meat production.

## Introduction

Cultured meat (also called cultivated meat or cell-based meat) is an emerging technology with the potential to overcome conventional animal agriculture’s limitations in terms of sustainability, animal welfare, and human health^1^. Briefly, cultured meat is defined as meat produced through cell culture and tissue engineering techniques, rather than by raising and slaughtering livestock. The potential benefits of cultured meat have been projected and discussed in prospective analyses and reviews^2–4^; however, substantial research and development is still required to make this technology economically viable at a scale and price-point comparable to conventional meat products^5^. Here, a key area of research continues to be the culture media, or the liquid substrates that provide nutrients, buffering, and functional factors (e.g., signaling factors or carrying proteins) to cells *in vitro* in order to promote cell expansion and maturation into relevant tissue types (typically muscle and fat). Specific challenges to media development for cultured meat are a traditional dependence on animal-derived components such as fetal bovine serum (FBS), or else the prohibitively expensive cost of serum-free media (SFM) formulations. As media are projected to account for over 90% of the production costs of cultured meat^6^, lowering the cost of appropriate animal component-free media is essential for achieving price parity with conventional meats.

Recently, we developed an animal-component free medium that was suitable for the expansion of bovine satellite cells (BSCs)^7^. This medium, termed Beefy-9, was developed by adding recombinant albumin to a previously developed medium (B8), which had itself been established for culturing induced pluripotent stem cells (iPSCs)^8^. While the addition of recombinant albumin rendered this medium suitable for BSC expansion, it also added substantial cost. Specifically, depending on its concentration, recombinant albumin addition resulted in a 50-400% increase in cost compared with standard B8. This price-dominance of albumin has been noted by others as well, including in a recent techno-economic analysis generated using data and input from current cultured meat companies^9^. As such, it is clear that cheaper alternatives must be developed in order to bring down costs of serum-free cell culture.

One promising option toward this goal is to use proteins extracted from low-cost and abundant sources. For instance, hydrolyzed proteins from rapeseed, wheat, soy, chickpeas, and other plants or agricultural byproducts have previously been used as albumin alternatives for culturing Chinese hamster ovary (CHO) and other mammalian cell types^10–15^. These sources offer the substantial benefits of being low-cost and highly scalable, and they can be sourced as the byproducts of other agricultural processes (such as the production of canola oil in the case of rapeseed protein). However, while the hydrolysis used in these studies can enable the production and purification of well-defined peptide fractions with specific functionalities in cell culture, it can also add processing complexities (e.g., hydrolytic enzymes or chemical treatments) which may impact the cost and scalability of these inputs in cultured meat applications. Thus, it may be valuable to use bulk plant protein extracts (i.e., nonhydrolyzed) in order to maximize process simplicity, affordability, and scalability.

The current work demonstrates that bulk protein extracts from oilseed protein meals can serve as albumin alternatives in Beefy-9. Specifically, rapeseed protein isolate (RPI) produced through simple alkali extraction and isoelectric precipitation can fully replace recombinant albumin in Beefy-9 culture of BSCs over short- and long-term culture, thereby dramatically lowering the cost of the cell culture medium. More, this modified serum-free medium (termed Beefy-R) supports enhanced cell growth compared with Beefy-9 while maintaining the myogenic phenotype and differentiation capacity of cells at least as well as Beefy-9. Together, these findings demonstrate the utility of bulk oilseed protein isolates (OPIs) and particularly RPI in replacing recombinant albumin in cell culture, thereby enabling substantial cost reduction for cultured meat production.

## Results

### Alkali extraction of protein meals enables simple and low-cost generation of protein isolates

Oilseed protein isolates (OPIs) from four plant sources were used in this study. These included Inca peanut (*Plukenetia volubilis)*, soybean (*Glycine max*), rapeseed (*Brassica napus*), and cottonseed (*Gossypium hirsutum*). OPIs were prepared using the same methods for each plant source. These involved alkali extraction (pH 12.5), isoelectric precipitation (pH 4.5), centrifugation, filtration, and ultrafiltration to concentrate the final protein solutions to 50 mg/mL (Fig. 1a). This method resulted in Inca peanut protein isolate (IPPI), soy protein isolate (SPI), rapeseed protein isolate (RPI), and cottonseed protein isolate (CPI). The isolates were either clear in color (IPPI and SPI) or reddish brown (RPI and CPI), and were slightly viscous. All OPIs could be snap-frozen, stored at −80°C, and thawed to room temperature without any impact on solubility, though long-term storage at 4°C or lyophilization resulted in aggregation and reduced solubility.

**Figure 1:**
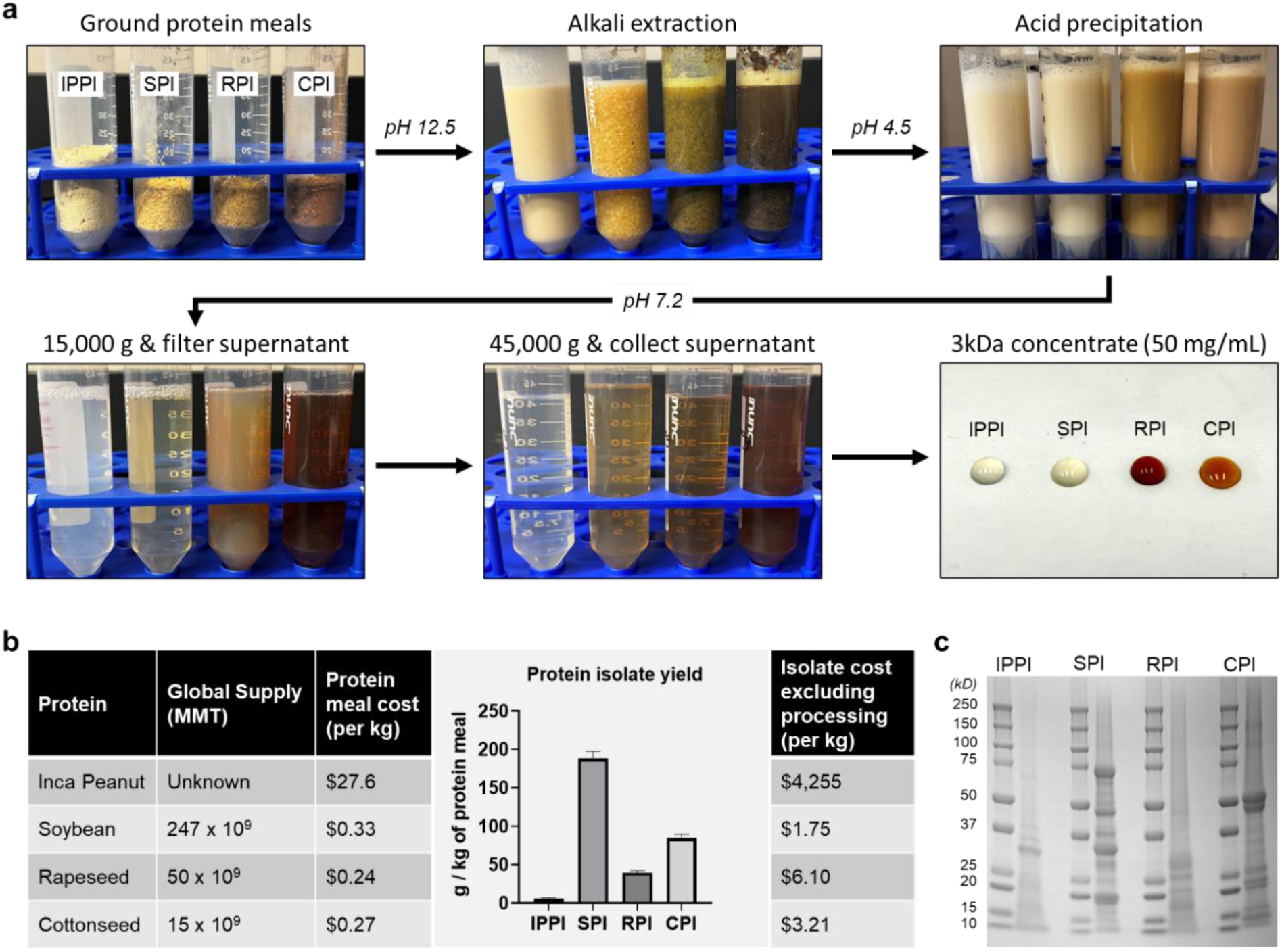
Protein isolate generation. **(a)** The methods used for generating OPIs involved alkali extraction, isoelectric precipitation, centrifugation, and filtration in order to generate concentrated protein solutions (50 mg/mL). The resulting OPIs were clear or reddish-brown in color, and slightly viscous. **(b)** Comparisons of global protein meal supply and cost, unoptimized protein yields, and OPI cost based on yields and starting protein meal cost (excluding processing costs from inputs such as chemicals, energy costs, filters, etc.). The results showed substantially reduced cost for SPI, RPI and CPI compared to IPPI. **(c)** SDS-PAGE of OPIs next to corresponding ladder for each protein. The results revealed heterogeneous mixtures of proteins <75 kDa, with larger average protein fractions for SPI and CPI compared with IPPI and RPI.

While the Inca peanut protein powder was originally produced for human consumption and was therefore relatively expensive ($27.6/kg), the three other oilseed protein meals were the byproduct of food oil production and are therefore attainable at low cost (less than $0.4/kg). The annual global supply of soybean meal, rapeseed meal, and cottonseed meal is 247, 50, and 15 trillion metric tons, respectively^16^. Following extraction and concentration, OPI yields were 6.5 g/kg, 188 g/kg, 39 g/kg, and 85 g/kg for IPPI, SPI, RPI, and CPI, respectively (Fig 1b). Using these yields and available data on the costs for protein meals, the costs of OPI starting materials were $4,255/kg, $1.75/kg, $6.10/kg, and $3.21/kg, respectively. These calculations do not include processing costs (e.g., solutions, electricity, filters, etc.), and so real-world cost-of-goods would likely be higher; however, even accounting for added processing expenses, the costs of SPI, RPI, and CPI are orders of magnitude lower than the cost of recombinant albumin available today, which is around $45,000/kg (Cellastim S, InVitria). This suggests that OPIs could be affordable albumin alternatives in cell culture. More, there is likely room for process optimization to increase yields, which was not explored in this study, and which would reduce costs further.

Following protein isolation, basic characterization was performed via SDS-PAGE (Fig 1c). The resulting gels revealed that OPIs were heterogeneous mixtures of proteins which were <75 kDa for all four plant sources. SPI and CPI showed higher levels of large proteins (>37 kDa), while the majority of proteins in IPPI and RPI were <37 kDa. Observed proteins likely correspond with fractions of various albumins and globulins, which are the largest fractions of seed storage proteins, and many of which exist in the size ranges observed for OPIs^17–22^. Ultimately, these results show that despite the use of the same extraction method for each plant source, the resulting protein mixtures varied substantially, suggesting that functionality may differ between OPIs produced through the same method.

### Oilseed protein isolates can replace albumin functionality in serum-free media for short-term growth

The primary bovine satellite cells (BSCs) used for this work have been extensively characterized and validated in previous studies^7,23^. To test the ability of OPIs to replace recombinant albumin in serum-free media, BSCs were cultured for four days in either B8 medium without supplementation, B8 medium supplemented with OPIs at a range of concentrations, or Beefy-9 medium (B8 + 0.8 mg/mL recombinant albumin). The results showed that over four days, SPI, CPI, and RPI supplementation significantly improved cell growth compared with B8 alone, while IPPI did not (Fig. 2a). Here, both SPI and RPI performed better than CPI, while RPI at 0.4 mg/mL performed the best of all supplements and completely recovered the efficacy of recombinant albumin (1.15-fold efficacy compared with Beefy-9). From these results, a new medium termed Beefy-R was developed which comprises B8 supplemented with 0.4 mg/mL RPI, and which suitably replaces Beefy-9 over short term culture. Brightfield microscopy of cells cultured in Beefy-R showed no morphological differences to those cultured in Beefy-9 (Fig. 1b). The final composition of Beefy-R is DMEM/F12 with L-ascorbic acid 2-phosphate (200 μg/mL), insulin (20 μg/mL), transferrin (20 μg/mL), sodium selenite (20 ng/mL), fibroblast growth factor-2 (FGF2-G3; 40 ng/mL), neuregulin 1 (NRG1; 0.1 ng/mL),transforming growth factor beta-3 (TGFβ; 0.1 ng/mL), and RPI (0.4 mg/mL). Methods for preparation are given in supplementary materials.

**Figure 2:**
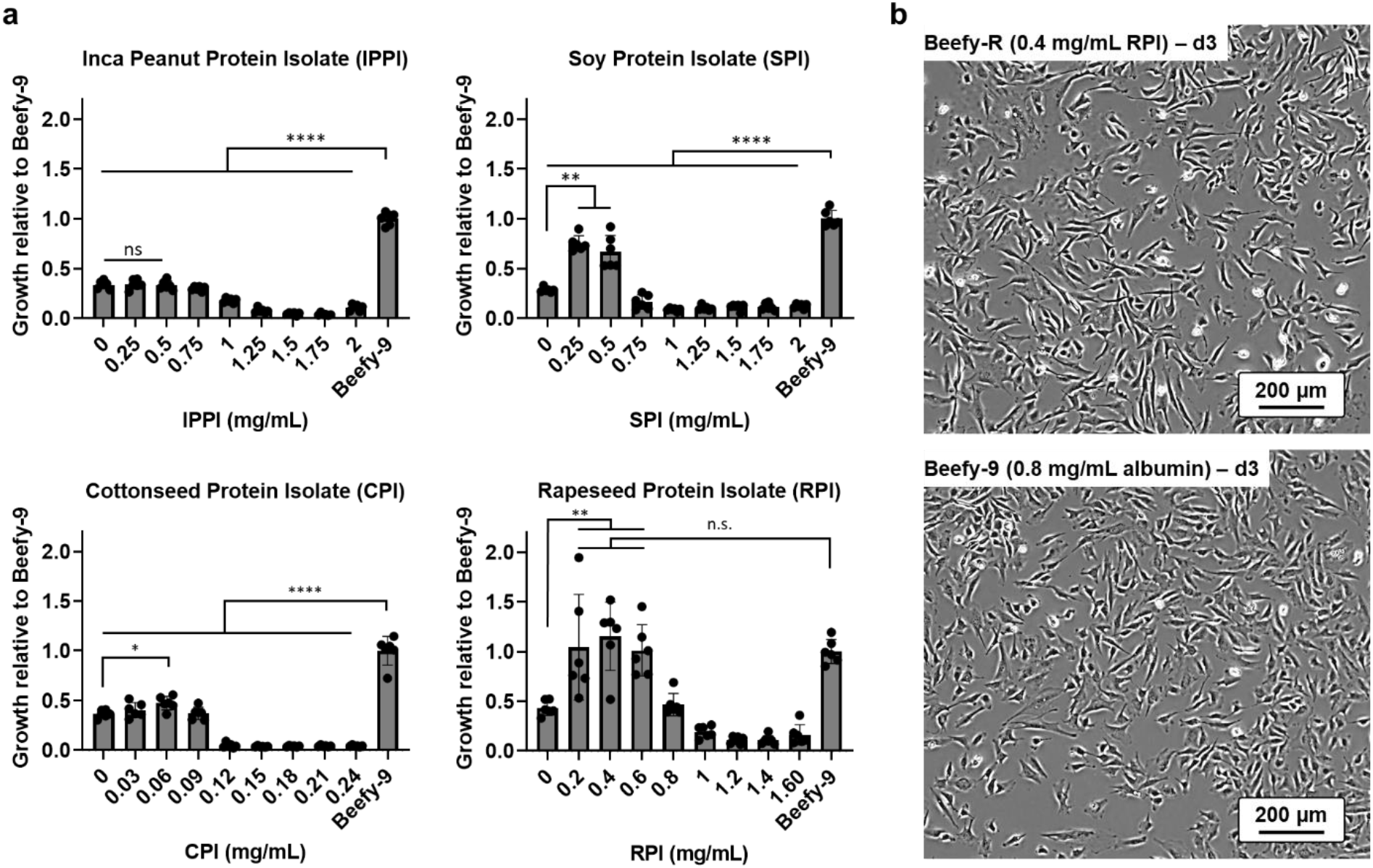
Short-term growth in OPI-supplemented serum-free media. **(a)** Four-day cell growth in B8 medium supplemented with various OPIs compared with B8 supplemented with 0.8 mg/mL recombinant albumin (Beefy-9). The results showed that SPI, RPI, and CPI significantly improved cell growth compared to no supplementation, but only RPI was able to recover cell growth comparable to Beefy-9 (1.15 times the growth of Beefy-9, not significant). *n* = 6 distinct samples; statistical significance was calculated by one-way ANOVA and is indicated for *p* < 0.05 (*), *p* < 0.01 (**), *p* < 0.001 (***), *p* < 0.0001 (****). **(b)** Brightfield images of BSCs on day three of growth in Beef-R or Beefy-9 showed no difference in cell morphology between conditions. Scale bars are 200 μm.

To briefly explore how protein extraction might impact RPI’s efficacy, protein isolates were prepared using slightly altered methods, including pre-extraction defatting with hexane, increasing salt concentration with sodium chloride, and extracting proteins with or without an overnight incubation at 4°C before final concentration and filtration steps. While hexane defatting reduced the optimum concentration of RPI to 0.2 mg/mL without any loss of efficacy, no significant difference was seen between optimum concentrations of the various extraction methods (Fig. S1). Thus, hexane defatting was avoided in generating Beefy-R due to the potential negative environmental impacts associated with this solvent. Additionally, to compare the efficacy of RPI with other potential albumin-replacing supplements, these same short-term experiments were performed using B8 supplemented with candidates including cyclodextrins^24^, off-the-shelf plant protein hydrolysates^25^, and molecular crowding agents^26^. However, none of these supplements were able to recover the efficacy of recombinant albumin in culture (Fig. S2), suggesting the unique potential of RPI in serum-free culture of bovine muscle cells.

### Beefy-R supports enhanced multi-passage expansion of BSCs compared with Beefy-9

Following short-term validation of RPI as an albumin alternative in serum-free media, multi-passage growth was assessed for BSCs in Beefy-R, Beefy-9, and a serum-containing growth medium (BSC-GM) (Fig. 3). For BSC-GM, tissue culture plates were coated with 0.25 μg/cm^2^ of recombinant laminin, while for Beefy-9 and Beefy-R, plates were coated with 1.5 μg/ cm^2^ of recombinant vitronectin, as determined during the development of Beefy-9. Similarly, as determined during the development of Beefy-9, recombinant albumin and RPI were withheld from the media while seeding for each passage, and were added only after cells had been allowed to attach to the culture vessels overnight^7^. The results showed that Beefy-R offered improved growth compared with Beefy-9 over four passages (Fig. 3a). Indeed, cells grown in Beefy-R achieved 11.7 population doublings over thirteen days compared with 10.6 doublings for cells growth in Beefy-9. Thus, twice as many cells were produced over two weeks when recombinant albumin was replaced with RPI. However, neither serum-free media formulation promoted growth equivalent to 20% FBS (14.8 population doublings), suggesting that further optimization is required.

**Figure 3:**
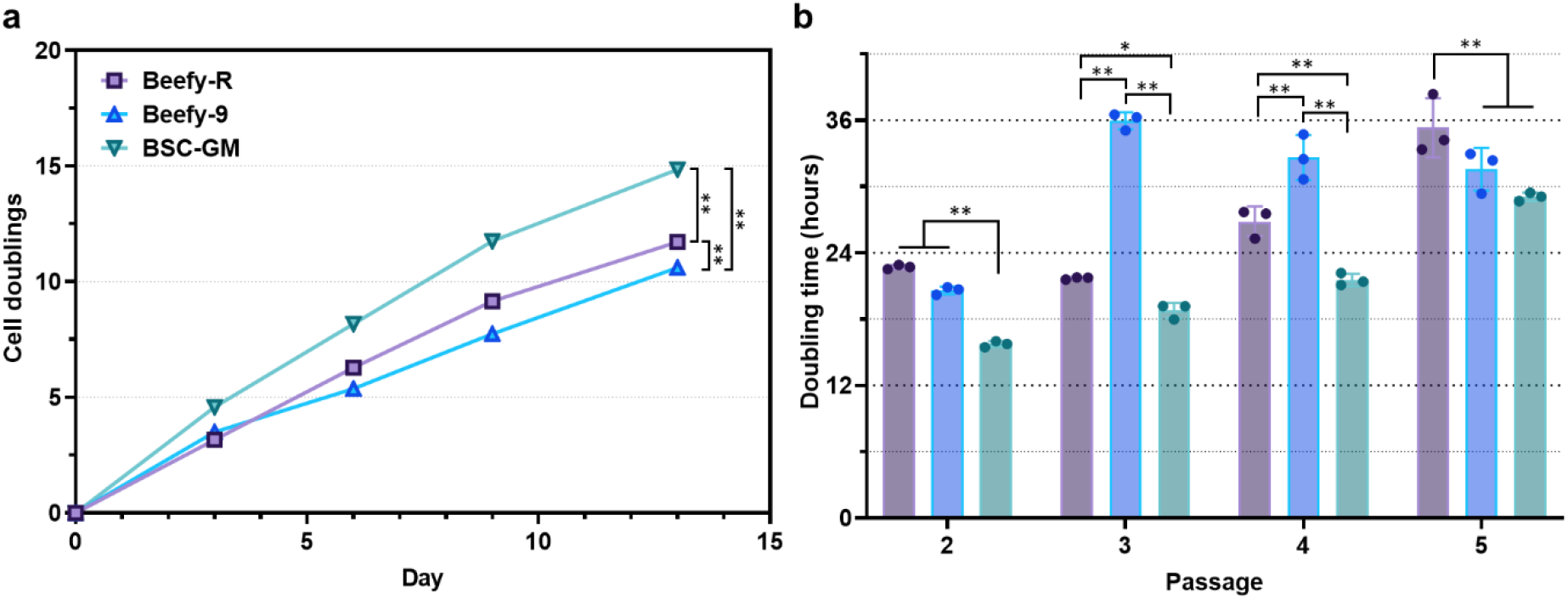
Multi-passage growth in Beefy-R. **(a)** Four-passage cell growth in various media. The results showed that Beefy-R improved growth over Beefy-9, though not to the degree of BSC-GM. *n* = 3 distinct samples; statistical significance was calculated by two-way ANOVA and is indicated for the final timepoint with *p* < 0.05 (*), *p* < 0.01 (**), *p* < 0.001 (***), *p* < 0.0001 (****). **(b)** Doubling time calculations for cell growth in (a). While growth slowed for all media types, results showed that Beefy-R maintained doubling times <24 hours for two passages, compared with one for Beefy-9 and three for BSC-GM. *n* = 3 distinct samples; statistical significance was calculated by two-way ANOVA and is indicated for *p* < 0.05 (*), *p* < 0.01 (**), *p* < 0.001 (***), *p* < 0.0001 (****).

Comparing growth rates over four passages, the average population doubling times for Beefy-R, Beefy-9, and BSC-GM were 26.6, 30.2, and 21.3 hours, respectively. While all media types showed an increase in doubling time with increased passage number, Beefy-R resulted in a slower deterioration of doubling rate than Beefy-9, though faster deterioration than BSC-GM. Together, these results showed that Beefy-R was able to improve BSC growth over Beefy-9, though not to the degree of BSC-GM. This suggests that RPI is able to not only replace albumin in serum-free medium, but indeed offers improved functionality.

### Beefy-R maintains BSC identity and myogenicity during culture

BSCs cultured for two passages in various media were then subjected to a range of analyses. These were performed on proliferating cells (70% confluency) and cells that had been cultured to confluency and treated with a previously described serum-free differentiation medium for two days.^27^ The same serum-free differentiation medium was used for cells cultured in both serum-free and serum-containing media, in order to keep differentiation conditions consistent. For the first analysis, RNA was harvested from cells and quantitative PCR (qPCR) was performed to assess expression of four genes: early satellite cell marker Paired-box 3 (Pax3), myogenic commitment marker Myoblast Determination Protein 1 (MyoD), early differentiation marker Myogenin, and terminal fusion and differentiation marker Myosin Heavy Chain (MHC). Data was analyzed as fold-change compared with BSC-GM proliferating cells (Fig 4a). For proliferating cells, the most notable differences between serum-free and serum-containing conditions existed for MyoD and Myogenin. Specifically, BSC-GM cells showed a significant (∼10-fold) increase in MyoD compared with Beefy-R or Beefy-9, and a significant decrease in Myogenin (∼6-fold). These results suggest that serum-containing BSC-GM was able to maintain an earlier muscle phenotype than serum-free Beefy-R or Beefy-9. For differentiating cells, the most notable differences existed for Myogenin and MHC, where both serum-free media showed a significant (∼3.5-fold) increase in Myogenin compared with BSC-GM, and Beefy-R showed a significant (∼3-fold) increase in MHC compared with BSC-GM. These results suggest enhanced differentiation of cells cultured in serum-free media, which is in line with the later-stage proliferative phenotype of these cells.

**Figure 4:**
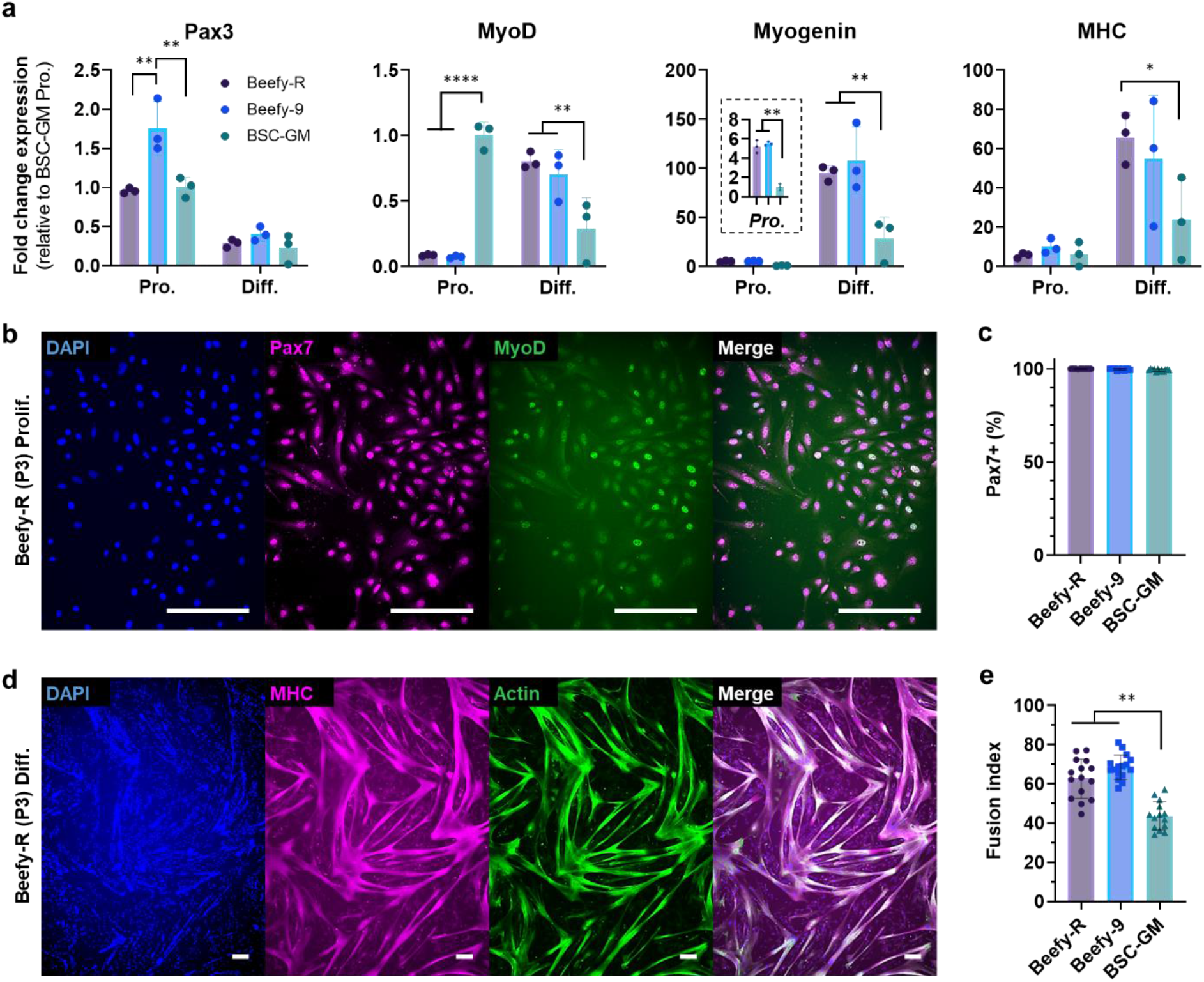
Phenotypic analysis of cells cultured in various media. **(a)** qPCR of proliferative cells (Pro.) cultured in various media, and cells following two days of differentiation (Diff.). In proliferating cells, results showed increased MyoD expression for BSC-GM and increased Myogenin expression for Beefy-R and Beefy-9. In differentiated cells, results showed increased MyoD, Myogenin, and MHC expression in Beefy-R and Beefy-9 compared with BSC-GM. *n* = 3 distinct samples; statistical significance was calculated by two-way ANOVA (or one-way ANOVA for Myogenin Pro. insert) and is indicated for *p* < 0.05 (*), *p* < 0.01 (**), *p* < 0.001 (***), *p* < 0.0001 (****). **(b)** Immunostaining of proliferative cells in Beefy-R for nuclei (DAPI, blue), Pax7 (magenta), and MyoD (green). The results showed ubiquitous staining for Pax7 and heterogeneous staining for MyoD. Similar staining for Beefy-9 and BSC-GM is given in Fig. S3. Scale bars are 200 μm. **(c)** Pax7 quantification revealed >99% Pax7+ for all three media types (Beefy-R = 99.98%; Beefy-9 = 99.70%; BSC-GM = 99.34%. *n* = 12 (4 images each for 3 distinct samples). **(d)** Immunostaining of differentiated cells from Beefy-R for nuclei (DAPI), MHC (magenta), and actin (green). Results showed robust formation of MHC-positive multinucleated myotubes. Similar staining for Beefy-9 and BSC-GM is given in Fig. S5. Scale bars are 200 μm. **(e)** Fusion index analysis of differentiated myotubes revealed significantly enhanced differentiation for Beefy-R and Beefy-9 compared with BSC-GM. *n* = 13-15 images for one distinct sample; statistical significance was calculated by two-way ANOVA and is indicated for *p* < 0.01 (**).

Along with qPCR, immunostaining for Pax7, MyoD, and MHC was performed on proliferative and differentiated cells. Proliferative cells showed ubiquitous expression of Pax7 and heterogeneous expression of MyoD for all media types (Fig. 4b & S3). Image analysis of Pax7 staining revealed that >99% of cells were positive for Pax7 for all three media types, suggesting that all three media maintained satellite cell identity to a high degree (Fig. 4c & S4). Differentiated cells showed myotube fusion and MHC staining for all media types, indicating robust differentiation following culture in both serum-free and serum-containing media (Fig. 4d & S5). Image analysis of MHC staining was performed to assess fusion index, or the percentage of nuclei that are present in MHC-positive myotubes (Fig. S6). The results showed that fusion index was significantly higher for Beefy-R and Beefy-9 compared with BSC-GM (Fig. 4e), which supports the qPCR results showing enhanced differentiation of cells cultured in serum-free media. Finally, brightfield images of passage 5 cells showed reduced lipid droplet formation in both Beefy-R and BSC-GM compared with Beefy-9 (Fig. S7). This is notable, as aberrant lipid accumulation was observed during the development of Beefy-9, and so replacement of albumin with RPI could reduce this phenotypic change when culturing BSCs in serum-free media^7^.

Together, qPCR and immunostaining data revealed that RPI maintained myogenic phenotype at least as well as recombinant albumin, but that both Beefy-R and Beefy-9 resulted in a later-stage phenotype than serum-containing media. This difference resulted in enhanced differentiation compared with BSC-GM, although this could also induce reduced stemness and proliferative capacity over longer-term culture. Ultimately, these results further support the utility of RPI for replacing recombinant albumin in serum-free culture and suggest that further Beefy-R media optimization is required to match the efficacy of BSC-GM.

### Proteomic analysis reveals differences between OPI composition

Proteomic analysis was performed on those OPIs that improved BSC growth (i.e., CPI, SPI and RPI) and Gene Ontology (GO) analysis was used to compare between samples. Specifically, mass-spectrometry data was gathered and compared against reference proteomes for rapeseed, cottonseed, and soybean. Identified proteins were weighted by spectral counts (SpC) and categorized by biological process, cellular component, and molecular function. The results showed that, while all OPIs differed from each other, SPI and RPI were more similar to each other than to CPI. For instance, looking at the biological process categorization (Fig. 5a), SPI and RPI had a substantially higher prevalence of proteins involved in “cellular process,” “metabolic process,” and “response to stimulus” processes, while CPI had a substantially higher prevalence of those involved in “developmental process,” “multicellular organismal process,” “reproduction,” and “reproductive process” processes. Looking at cellular component categorization (Fig. 5b), SPI and RPI had substantially higher prevalence of cytoplasmic proteins, while CPI had a substantially higher prevalence of membrane proteins. SPI also had a substantially higher prevalence of ribosomal proteins, and all three OPIs had a high prevalence of proteins located in intracellular organelles. Finally, looking at molecular function categorization (Fig. 5c), SPI and RPI had a higher prevalence of “binding” and “catalytic activity” proteins, while CPI had a higher prevalence of “nutrient reservoir activity” proteins. It is possible that the enhanced efficacy of SPI and RPI compared with CPI in Fig. 2 was attributable to these trends in protein composition, and so certain protein categories that are more prevalent in SPI and RPI could be the functional components that are responsible for their efficacy in serum-free media.

**Figure 5:**
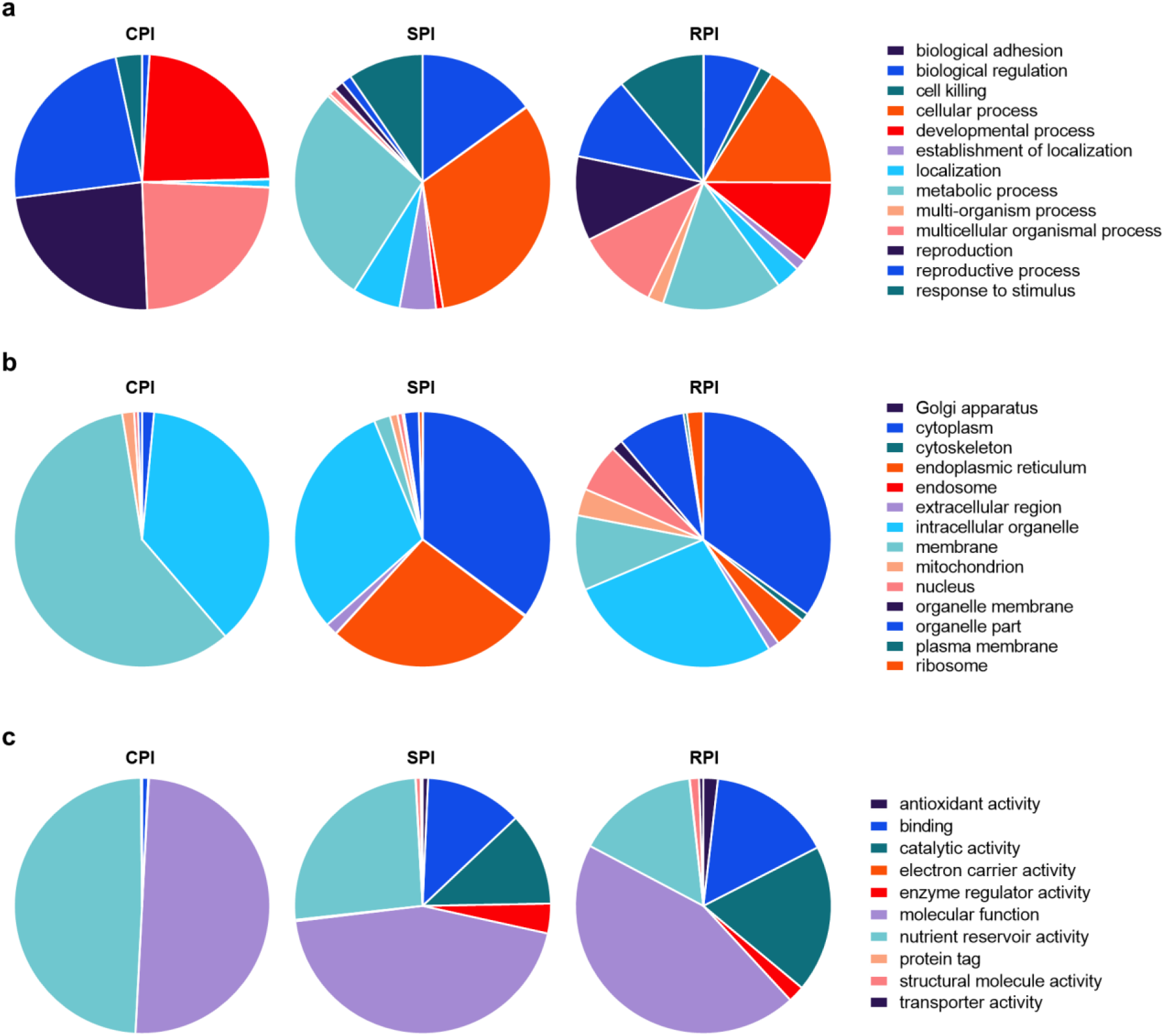
Gene ontology (GO) classification of proteins in OPIs. **(a)** GO terms of proteins classified under “Biological Process” for three OPIs. The results showed increased prevalence of “cellular process” and “metabolic process” proteins in SPI and RPI compared with CPI, and increased concentrations of “developmental process,” “multicellular organismal process,” “reproduction,” and “reproductive process” proteins in CPI. Frequency of protein types was weighted by spectral count. **(b)** GO terms of proteins classified under “Cellular Component” for three OPIs. The results showed increased prevalence of “cytoplasm” proteins for SPI and RPI compared with CPI, increased concentrations of “endoplasmic reticulum” proteins for SPI, and increased concentrations of “membrane” proteins for CPI. Frequency of protein types was weighted by spectral count. **(c)** GO terms of proteins classified under “Molecular Function” for three OPIs. The results showed increased concentrations of “binding” and “catalytic activity” proteins for SPI and RPI compared with CPI, and increased concentrations of “nutrient reservoir activity” proteins for CPI. Frequency of protein types was weighted by spectral count.

Looking at specific proteins, the three most prevalent for RPI included one protein of the 2s seed storage albumin family (napin; UniProt accession #A0A078IPA5) and two proteins of the 11s globulin family (cupins; UniProt accession #A0A078HBQ3 and # A0A078G0Q2). These each comprised ∼4% of total SpCs in RPI. The three most prevalent proteins for SPI included three cupin type-1 11s globulins comprising 9%, 7%, and 7% of total SpCs (UniProt accession #P0DO16, #P11827, and #P02858, respectively). Lastly, the three most prevalent proteins for CPI included one cupin type-1 7s globulin comprising 44% of total SpCs (UniProt accession #A0A1U8LMX7), and two 11s globulins comprising 21%, and 17% of total SpCs (UniProt accession #A0A1U8KKK7 and # A0A1U8KAE1), respectively. These results correspond with SDS-Page results in Fig. 1, the molecular weights of which suggested predominantly albumins and globulins for the OPIs.

Interestingly, the highest concentration 2s albumin storage protein in SPI comprised 1.2% of total SpCs, and the highest concentration 2s albumin in CPI comprised 1% of total SpCs. This was in comparison with 3.7% for RPI. This increased concentration of 2s albumin could have had some impact on the efficacy of RPI compared with SPI and CPI as a serum albumin replacement, though further research is required to parse out the functional contribution of specific proteins. For instance, RPI also had a substantially higher prevalence of oil body associated proteins (OBAPs) than SPI and CPI, and so it is impossible to definitively attribute RPI’s efficacy to its albumin fraction without further studies. Further, while spectral counting is typically considered a suitable quasi-quantitative metric for analyzing protein prevalence, confounding factors such as protein size and instrument differences can limit the quantitative value of these analyses^28^. As such, further quantitative proteomic analyses should be performed to better understand OPI profiles in order to elucidate functional differences.

## Discussion

Replacing recombinant albumin in serum-free media is a key priority for cultured meat research and development. Specifically, due to the high concentrations of albumin required in relevant serum-free media to-date, it is expected that recombinant techniques—even at their most efficient—will be unable to provide sufficiently affordable albumin to enable scalable, low-cost bioprocesses^29^. It has therefore been suggested that functional albumin alternatives are necessary to achieve economic viability at scale. These alternatives should be effective, low-cost, scalable, food-safe, and sustainable. Agricultural byproducts offer many advantages towards achieving these goals, including the valorization of high-volume waste streams and availability of low-cost inputs which can be processed in a simple, food safe manner. In this work, an effective serum-free medium (termed Beefy-R) was developed as validation of the efficacy of non-hydrolyzed oilseed protein isolates (particularly rapeseed protein isolate) as effective albumin alternatives. RPI is scalable to produce, relying on only simple and low-cost alkali/acid extraction and filtration steps. RPI was also shown to exceed recombinant albumin in promoting BSC growth while maintaining myogenicity.

While determining total cost-of-goods of RPI at scale (including processing costs such as electricity, filtration, chemicals, and waste management) is outside the scope of this work, the low cost of input materials and simplicity/scalability of extraction and post-processing should result in a highly affordable media input. Specifically, the input cost of rapeseed protein meal to make enough RPI for one liter of Beefy-R is $0.002. This results in a total media cost of $49.7/L for Beefy-R, compared with $74.3/L for Beefy-9, and $184.2/L for BSC-GM, when components are bought in bulk (Fig. 6).

**Figure 6:**
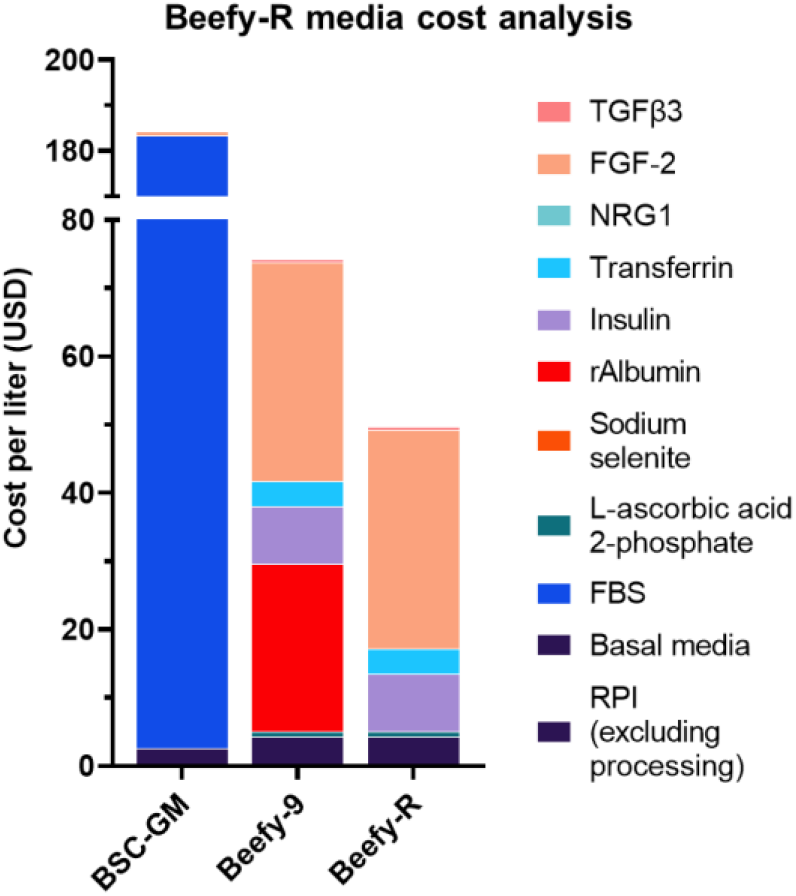
Media cost analysis. When purchasing components in bulk, costs of media in this study are $184.2/L for BSC-GM, $74.3/L for Beefy-9, and $49.7/L for Beefy-R (excluding RPI processing costs).

After overcoming the high cost of recombinant albumin with RPI, FGF-2 remains a key cost contributor in Beefy-R. Here, price is driven by the currently high cost of recombinant FGF-2, rather than the concentration used (40 ng/mL). As such, it is likely that process scale-up can drive down costs substantially by leveraging economies of scale. Methods to optimize and enhance recombinant growth factors or overcome their requirement in cell culture could also dramatically reduce these costs^8,30–33^. Additionally, during Beefy-9 development, it was shown that lowering the concentration of FGF-2 to 5 ng/mL did not drastically reduce growth, and so this simple strategy could reduce the cost of Beefy-R to $21.7/L. Similarly, recent studies have shown robust BSC growth in serum-free media containing reduced insulin (10 μg/mL compared with 20 μg/mL in Beefy-9), reduced FGF-2 (10 μg/mL compared with 40 μg/mL in Beefy-9), and reduced transferrin (5.5 μg/mL compared with 20 μg/mL in Beefy-9)^34^. Thus, reducing these components could be explored to optimize Beefy-R. Finally, it should be noted that the protein isolation process used in this work leaves behind a number of proteins that are insoluble at physiological pH, and which could themselves be useful byproducts for the production of resources such as scaffolds for cultured meat production processes. For instance, soy protein isolates have recently been used to form edible films or 3D scaffolds for culturing BSCs for cultured meat^35,36^. It is possible that similar scaffolds could be generated using the rapeseed protein fraction that remains after the final centrifugation steps used in this study, thereby further reducing the cost of RPI by valorizing the unused portion.

From a consumer and regulatory standpoint, several features of RPI should be considered and studied prior to implementation in cultured meat development. Specifically, the food-safety of RPI must be validated as appropriate for cultured meat production processes^37^. This is important, as rapeseed protein meal is typically used as animal feed or fertilizer, and so does not necessarily have the same quality control requirements that should be established for cultured meat production. Possible hazards include contaminating microorganisms in the protein meal that must be removed in order to use RPI for culture media. In this study, filtration through 0.45-micron filters was used to sterilize RPI; however, other sterilization methods might be required at scale to balance scale and efficacy. Additionally, oilseed proteins can be allergenic for some consumers, including 11s globulins prevalent in OPIs^38^. Further characterization should therefore be performed to determine the potential allergenicity of OPIs, as well as whether or not any allergens are incorporated into a final cultured meat product. Finally, the sustainability and scalability of RPI and other OPIs should be validated through dedicated techno-economic and life-cycle analyses.

Lastly, while this work confirms the utility of RPI as an albumin alternative, substantial future work should be pursued in order to further explore OPIs and other plant protein extracts as albumin alternatives. This could include fractionation of OPIs to identify functional constituents, optimizing extraction methods to maximize yields and efficacy, exploring other oilseeds and agricultural waste streams for functional components, and optimizing other media components for enhanced cell growth and differentiation^34,39^. Here, computational methods can be leveraged to identify additional low-cost inputs (e.g., functional protein fractions) and to optimize media formulations^40,41^. Finally, future work could explore plant protein extracts as alternatives for other costly recombinant proteins, such as insulin and transferrin, and could develop media for other cell types, such as preadipocytes and iPSCs.

In summary, the present work validates the utility of OPIs, particularly RPI, in replacing recombinant albumin in serum-free media for BSCs. The resulting media, Beefy-R, enhanced cell growth compared with albumin-containing Beefy-9 while reducing costs and maintaining satellite cell phenotype and myogenicity. Protein analysis of OPIs revealed compositional differences which likely correlate with functional disparities. While substantial opportunities for further research and optimization exist, this work addresses a key challenge facing cultured meat development and lays a foundational platform for achieving truly cost-effective serum-free media formulations.

## Materials and Methods

### Bovine satellite cell culture and routine maintenance

Primary BSCs used in this study were isolated and characterized previously using tissue from a 2-week old Simmental calf^7^. For routine cell maintenance, cells were cultured in 37°C with 5% CO_2_ on tissue-culture plastic coated with 0.25 ug/cm^2^ iMatrix recombinant laminin-511 (Iwai North America #N892021, San Carlos, CA, USA). BSC growth media (BSC-GM) was used, comprised of DMEM+Glutamax (ThermoFisher #10566024, Waltham, MA, USA), 20% fetal bovine serum (FBS; ThermoFisher #26140079), 1 ng/mL human FGF-2 (ThermoFisher #68-8785-63), and 1% antibiotic-antimycotic (ThermoFisher #1540062). For routine culture, cells were expanded to a maximum of 70% confluence, harvested using 0.25% trypsin-EDTA (ThermoFisher #25200056), counted using an NC-200 automated cell counter (Chemometec, Allerod, Denmark) and either seeded at a density of 2,000-5,000 cells/cm^2^ on new tissue culture plastic with growth medium and laminin, or else frozen in FBS with 10% Dimethyl sulfoxide (DMSO, Sigma #D2650, St. Louis, MO, USA).

### Oilseed protein isolate generation

OPIs were generated from protein meals from four plant sources. These were Inca peanut (*Plukenetia volubilis)*, which was obtained as a protein powder (Imlak’esh Organics #765857605340, Santa Barbara, CA, USA); soybean (*Glycine max*), which was obtained as ground protein meal (Roots organics #715110, Eugene, OR, USA); rapeseed (*Brassica napus*), which was obtained as protein cakes (Joshua Roth Limited #6046, Albany, OR, USA); and cottonseed (*Gossypium hirsutum*), which was obtained as ground protein meal (Down To Earth #07859, Eugene, OR, USA). Rapeseed protein cakes were ground in a standard coffee grinder to generate a ground protein meal, after which protein isolation procedures were the same for all samples, based on adapted methods from several previously reported studies^10,18,42–48^. Briefly, for each protein, 10 g of protein meal was mixed with 100 mL of DI water, and the resulting slurry was adjusted to a pH of 12.5 using 5M NaOH. The slurry was mixed at room temperature in a beaker with a magnetic stir bar for one hour to extract proteins. Next, the mixture was centrifuged at 15,000 g and room temperature for 10 minutes to pellet non-soluble components, and the protein-containing supernatant was collected. This supernatant was adjusted to pH 4.5 with 6M HCl in order to precipitate out proteins, and solution was again centrifuged at 15,000 g and room temperature for 10 minutes. The supernatant was discarded, and the protein-containing pellet was resuspended in 100 mL of DI water which was adjusted to pH 12.5 using 5M NaOH. Proteins were encouraged to redissolve and redisperse in alkali solutions using a magnetic stir-bar and mechanical disruption of the pellet. When proteins were completely dispersed, pH was adjusted to 7.2 using 6N HCl in order to crash out those proteins that were not soluble at physiological pH. This mixture was centrifuged at 15,000 g and room temperature for 30 minutes, and the supernatant was filtered through a 10 μm filter before being centrifuged at 45,000 g and 4°C for three hours to completely pellet any suspended insoluble particles. The resulting OPI-containing supernatant was collected, filtered through an 0.22 μm filter, and concentrated with 3-kDa cutoff ultrafiltration tubes (Pall #MAP003C37, Port Washington, NY, USA) by centrifugation at 4,500 g and 4°C until 50- to 100-fold concentration had been achieved.

Concentrated proteins were filtered through 0.45 μm filters (Sigma #SLHV033R) and were then quantified using a Pierce BCA kit (ThermoFisher #23227) according to the manufacturers protocol. Once protein concentrations had been determined, they were adjusted to 50 mg/mL for all samples using sterile ultrapure water, and aliquots were snap-frozen in liquid nitrogen before moving them to long-term storage at −80°C. An easy-to-follow protocol for generating RPI is given in Supplementary Information.

To test how different protein isolation methods affected growth (Fig. S1), the following adjustments to the protocol were made: first, an overnight incubation at 4°C was added between the 10 μm filter step and the centrifugation at 45,000 g. Second, this same overnight incubation was added, but the protein solution was also supplemented with NaCl at a concentration of 120.6 mM. Third, hexane-defatting was performed before extraction by adding 10 g of protein meal to 100 mL of hexane (Sigma #270504-1L) and incubating the mixtures at room temperature on a stir-plate for one hour. After defatting, hexane was removed by decanting and protein meals were allowed left in the fume hood until fully dry. Defatted protein meals were then used for extraction as mentioned above, including the overnight incubation at 4°C between 10 μm filtration and 45,000 g centrifugation. As noted in the results, no significant differences were noticed between these changes, except that hexane defatted RPI showed optimal performance at a reduced concentration. It should also be noted that preliminary experiments for this study only used centrifugation up to 12,000 g and still saw RPI efficacy. It is therefore expected that lower-speed centrifugation will work in settings that do not have large, high-speed centrifuges, though times may need to be increased to achieve sufficient pelleting to avoid clogging filters.

### Oilseed protein isolate characterization: SDS-PAGE

For initial protein characterization, an SDS-Polyacrylamide gel (SDS-PAGE) was generated using the Laemmli protocol. Briefly, 120 μg of each OPI solution was diluted in 50 μL of water and mixed with 10 μL of 6x Laemmli reducing buffer (ThermoFisher #J61337-AC). Solutions were heated to 95°C for 10 minutes, centrifuged at 13,000 g for 5 minutes, and 10 μL of solution was loaded into a 4-20% Tris-glycine gel (ThermoFisher #XP04200PK2) along with 3 μL of Precision Plus protein ladder (Bio-Rad #1610374), and gels were run at 125 V for 1.25 hours in Tris-Glycine running buffer (ThermoFisher #LC2675).

### Short-term growth studies

Hi-Def B8 medium was prepared by adding Hi-Def B8 aliquots (Defined Bioscience #LSS-201, San Diego, CA, USA) to DMEM/F12 (ThermoFisher #11320033) along with 1% antibiotic-antimycotic. Beefy-9 medium was prepared by adding 0.8 mg/mL of recombinant albumin (Sigma #A9731-1G) to Hi-Def B8. Next, B8 medium supplemented with various concentrations of OPIs or other supplements (Fig. S2) were prepared. Cells were thawed into 96-well plates in BSC-GM with 0.25 μg/cm^2^ of recombinant laminin-511 and at a cellular density of 2,500 cells/cm^2^. Cells were allowed to adhere overnight in serum-containing media in order to ensure consistent starting cell numbers, after which cells were washed once with DPBS (ThermoFisher #14190250) and media was changed to either B8, B8 + supplements (e.g., OPIs), or Beefy-9. Two replicate plates were generated for each experiment, one of which was imaged, aspirated and frozen at −80°C on day three for dsDNA quantification, and one of which had its media refreshed on day three and was then aspirated and frozen at −80°C on day four for dsDNA quantification. This quantification was performed using a FluoReporter dsDNA quantification kit (ThermoFisher #F2962) according to the manufacturer’s instructions. Fluorescence was read on a Synergy H1 microplate reader (BioTek Instruments, Winooski, VT, USA) using excitation and emission filters centered at 360 and 490 nm.

### Multi-passage growth studies

Once short-term efficacy had been determined, multi-passage growth studies were performed to further validate the utility of RPI-supplemented Beefy-R medium. These studies used methods previously established in the development of Beefy-9^7^. Briefly, BSCs (passage 2) were seeded into triplicate wells of 6-well plates (Corning #353046, Corning, NY, USA) in BSC-GM with 0.25 ug/cm^2^ iMatrix laminin-511. After allowing cells to adhere overnight, cells were washed 1x with DPBS and fed either BSC-GM, Beefy-9, or Beefy-R (Hi-Def B8 supplemented with 0.4 mg/mL RPI). Beefy-9 and Beefy-R were prepared immediately before use. For passaging, cells were cultured to 70% confluency, rinsed 1x with DPBS, and dissociated with 500 μL of TrypLE Express (ThermoFisher #12604021). After incubating cells at 37°C for 10 minutes, plates were vigorously tapped to dislodge cells, and cells were collected with an additional 1.5 mL of either BSC-GM or Hi-Def B8, depending on whether they were the serum-containing or serum-free samples. Cells were then counted using an NC-3000 automated cell counter (Chemometec), centrifuged at 300 g for 5 minutes, resuspended in either BSC-GM or Hi-Def B8, re-counted, and seeded onto new 6-well plates at 2,500 cells/cm^2^ with 0.25 μg/cm^2^ of iMatrix laminin-511 for BSC-GM cells, or 1.5 μg/cm^2^ of truncated recombinant human vitronectin (Vtn-N; ThermoFisher #A14700) for serum-free cells. After allowing cells to adhere overnight, media was replaced with BSC-GM, Beefy-R, or Beefy-9, as appropriate. This process was repeated over 13 days and four passages (P2-P5) with cell feeding every two days. As before, Beefy-R and Beefy-9 were prepared immediately before use. When seeding passage three, additional wells were seeded for qPCR and immunostaining analyses.

### Serum-free differentiation

A population of cells at passage three was cultured to confluency in appropriate media, at which point media was changed to a previously described serum-free differentiation medium containing Neurobasal (Invitrogen #21103049, Carlsbad, CA, USA) and L15 (Invitrogen #11415064) media in a 1:1 ratio, supplemented with 10ng/mL insulin-like growth factor 1 (IGF-1; Shenandoah Biotechnology #100-34AF-100UG, Warminster, PA, USA), 100 ng/mL epidermal growth factor (EGF; Shenandoah Biotechnology #100-26-500UG), and 1% antibiotic-antimycotic ^49^. Cells were differentiated for 2 days before performing relevant analysis.

### Gene expression analysis

To assess relative gene expression between BSCs cultured in various media types, qPCR was performed on proliferative (70% confluency) and differentiated cells from P3 following standard protocols. Briefly, RNA was harvested using an RNEasy Mini kit (Qiagen #74104, Hilden, Germany) and cDNA was prepared via the iScript cDNA synthesis kit (Bio-Rad #1708890, Hercules, CA, USA) using 1 μg of RNA for each reaction. Next, qPCR was performed using 2 μL of cDNA and 1x TaqMan Fast Universal PCR Master Mix without AmpErase UNG (ThermoFisher #4352042). Primers used in this study were: 18S (ThermoFisher #Hs03003631), Pax3 (ThermoFisher #Bt04303789), MyoD1 (ThermoFisher #Bt03244740), Myogenin (ThermoFisher #Bt03258929), and Myosin Heavy Chain (ThermoFisher #Bt03273061). Reactions were performed according to the manufacturer’s instructions on a Bio-Rad CFX96 Real Time System thermocycler, and results were analyzed as 2^-ΔΔct^ normalized to expression of the 18S housekeeping gene and analyzed relative to proliferative BSC-GM expression for each gene.

### Immunostaining

Along with qPCR, immunocytochemistry was performed to assess phenotype of both proliferative and differentiated cells from each media. Cells were fixed with 4% paraformaldehyde (ThermoFisher #AAJ61899AK) for 30 minutes, washed 3x in DPBS, and permeabilized for 15 minutes with 0.5% Triton-X (Sigma # T8787) in DPBS. Cells were then rinsed 3x in PBS-T (DPBS containing 0.1% Tween-20 [Sigma #P1379]) and blocked for 45 minutes using a blocking buffer containing DPBS with 5% goat serum (ThermoFisher #16210064) 0.05% sodium azide (Sigma #S2002) before adding primary antibodies and incubating overnight at 4°C. For proliferative cells, primary antibodies used were for Pax7 (ThermoFisher #PA5-68506) diluted 1:500 in blocking buffer and MyoD (ThermoFisher #MA5-12902;1:100). For differentiated cells, primary antibodies used were for MHC (Developmental studies hybridoma bank #MF-20, Iowa City, IA, USA; 4 μg/mL) and Phalloidin-488 (Abcam #ab176753, Cambridge, UK; 1:1,000). Following primary antibody incubation, cells were rinsed 3x in PBS-T, incubated for 15 minutes at room temperature in blocking buffer, and treated with secondary antibodies in blocking buffer for one hour at room temperature. Secondary antibodies for proliferative cells were anti-rabbit (ThermoFisher #A-11072; 1:500) and anti-mouse (ThermoFisher #A-11001) for Pax7 and MyoD, respectively, as well as a DAPI nuclear stain (Abcam #ab104139, Cambridge, UK; 1:1,000). Secondary antibodies for differentiated cells were anti-mouse (ThermoFisher #A-11005) for MHC, as well as a DAPI nuclear stain. Finally, cells were rinsed 3x with DPBS before imaging.

Imaging was performed with a KEYENCE BZ-X810 fluorescent microscope (Osaka, Japan). For Pax7 quantification and fusion index analyses, batch images were taken with the 10x objective at random points of culture wells selected by the KEYENCE software. Images were analyzed using ImageJ software. Briefly, for Pax7 quantification, each nucleus in the DAPI channel was established as a discrete region of interest (ROI), and these ROIs were added to the Pax7 channel after thresholding at a consistent value. Percentage of ROIs containing Pax7 signal was recorded (Fig. S4). For fusion index quantification, total nuclei were counted in the DAPI channel, after which MHC channels were used to generate nuclear selections following the application of a consistent threshold. Nuclei within the selections were counted, and the ratios of selected nuclei to total nuclei were recorded as fusion indices.

### Proteomics

Proteomic analysis was performed at the Massachusetts Institute of Technology’s Koch Institute Proteomics core. First, Proteins were reduced with 10mM dithiothreitol (Sigma) for 10 minutes at 95°C and then alkylated with 20mM iodoacetamide (Sigma) for 30 minutes at 25°C in the dark. Proteins were than digested with trypsin on S-trap micro columns (Protifi #C02-micro-80) per the manufacturer’s instruction. Next, the tryptic peptides were separated by reverse phase HPLC (Thermo Ultimate 3000) using a Thermo PepMap RSLC C18 column (2um tip, 75umx50cm #ES903) over a 100-minute gradient before nanoelectrospray using an Exploris mass spectrometer (Thermo). Solvent A was 0.1% formic acid in water and solvent B was 0.1% formic acid in acetonitrile. The gradient conditions were 1% B (0-10 min at 300 nL/min), 1% B (10-15 min, 300 nL/min to 200 nL/min), 1-7% B (15-20 min, 200nL/min), 7-25% B (20-54.8 min, 200nL/min), 25-36 B (54.8-65 min, 200nL/min), 36-80% B (65-65.5 min, 200 nL/min), 80% B (65.5-70 min, 200nL/min), 80-1% B (70-70.1 min, 200nL/min), 1% B (70.1-90 min, 200nL/min). The mass spectrometer was operated in a data-dependent mode. The parameters for the full scan MS were: resolution of 120,000 across 375-1600 *m/z* and maximum IT 25 ms. The full MS scan was followed by MS/MS for as many precursor ions in a two second cycle with a NCE of 28, dynamic exclusion of 20 s and resolution of 30,000.

Raw mass spectral data files (.raw) were searched using Sequest HT in Proteome Discoverer (Thermo). Sequest search parameters were: 20 ppm mass tolerance for precursor ions; 0.05 Da for fragment ion mass tolerance; 2 missed cleavages of trypsin; fixed modification were carbamidomethylation of cysteine and TMT 10-plex modification on the lysines and peptide N-termini; variable modifications were methionine oxidation, tyrosine, serine and threonine phosphorylation, methionine loss at the N-terminus of the protein, acetylation of the N-terminus of the protein and also Met-loss plus acetylation of the protein N-terminus. For peptide groups data only PSMs with a Xcorr score greater than 2, isolation interference less than or equal to 30 and a deltaM(ppm) between −3 and 3 were used. For Gene Ontology (GO) analysis of the data, gene ontology annotation (GOA) files from UniProt were referenced in Scaffold (Proteome Software, Inc., Portland, OR; version 4.7.3). The GOA files used for CPI, SPI, and RPI were 4102840.G_hirsutum_Upland_cotton_Gossypium_mexicanum.goa, 23297.G_max.goa, and 388940.B_napus_Rape.goa, respectively. Weighted spectral counts of the GO terms in each of the three branches (molecular function, biological process, and cellular component) were calculated by summing all counts in each branch category, and were averaged between three technical replicates for each OPI.

### Statistical analysis

GraphPad Prism 9.0 software (San Diego, CA, USA) was used to perform all statistical tests. Specifically, for short-term growth studies, one-way ANOVA was performed along with a Tukey’s HSD post-hoc test between all samples, and statistical significance was indicated (Fig. 2a) for comparisons with Beefy-9 and with B8 (0 mg/mL OPI supplementation). For multi-passage analyses, two-way ANOVA was performed along with a Tukey’s HSD post-hoc test between all samples of a certain timepoint, and statistical significance was indicated for comparisons at day 13 only (Fig. 3a) or for comparisons at all timepoints (Fig. 3b). For qPCR studies, two-way ANOVA was performed with a Tukey’s HSD post-hoc test between all samples of a certain group (Pro. or Diff.), and statistical significance was indicated (Fig. 4a) for all comparisons. For fusion index studies, a one-way ANOVA was performed along with a Tukey’s HSD post-hoc test between samples (15 images per condition, except in the case where imaging artifacts clearly impacted image quality), and statistical significance was indicated (Fig. 4e) for all comparisons. P values <0.05 were treated as statistically significant, and all errors are given as ± standard deviation.

## Competing interests

The authors declare no competing interests.

## Author contributions

**AJS**: Conceptualization, Methodology, Investigation, Formal Analysis, Visualization, Writing-original draft preparation, Writing-reviewing and editing. **MLR:** Investigation. **MS:** Investigation. **ABM:** Formal Analysis. **MKS:** Methodology, Formal Analysis. **DLK:** Conceptualization, Resources, Writing-reviewing and editing, Supervision, Funding acquisition.

## Acknowledgements

We thank New Harvest for their support of this work. We also thank Dr. Cameron Semper for advice around protein isolation and protein chemistry, Dr. Eugene White for assistance with the satellite cell isolations that made this work possible, and Drs. Constancio Gonzalez Obeso and Ryan Scheel for their advice around protein extraction protocols. We thank the Koch Institute’s Robert A. Swanson (1969) Biotechnology Center for technical support, specifically the Proteomics Core and Richard Schiavoni. Thanks to Steven Rees and Defined Bioscience for their scientific input and generous contribution of reagents. This work was supported by the New Harvest Graduate Fellowship Program, the U.S. Department of Agriculture (2021-69012-35978), and the National Institutes of Health (P41EB002520).

## Data availability

The data supporting this study are available within the article’s Supplementary files. Extra data are available from the corresponding author upon request.

## Notes

### Competing Interest Statement

The authors have declared no competing interest.

