## Supplementary Materials for "A Beefy-R culture medium: replacing albumin with rapeseed protein isolates"

*Supplementary Figure 1: Comparing RPI extraction methods*

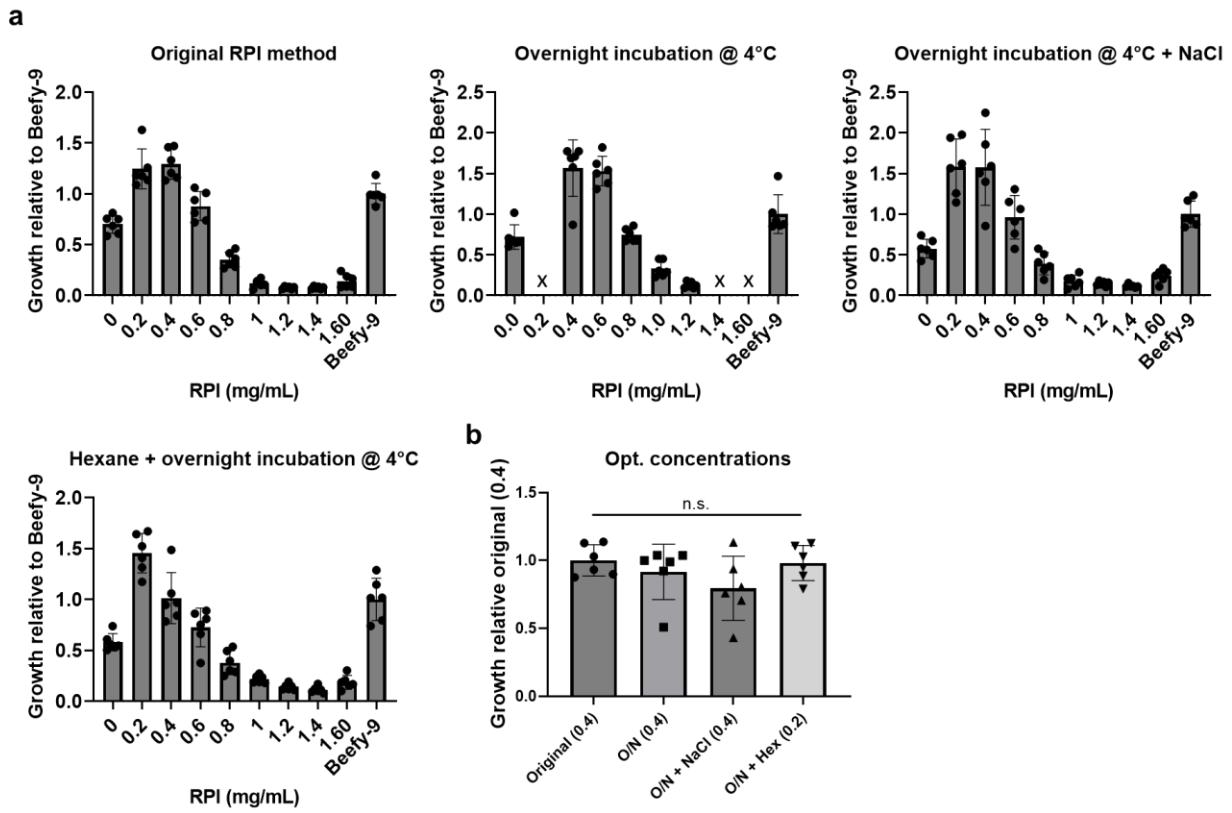


**Figure S1: Comparison of RPI extraction methods**. **(a)** RPI extracted using various methods (original described in *Materials and Methods*, original method plus an overnight incubation at 4°C, original method plus an overnight incubation at 4°C and with 120.6 mM NaCl added to the protein solution, original method plus an initial hexane defatting step for one hour and an overnight incubation at 4°C. Results showed no clear difference in cell growth of trends for the different extraction methods. **(b)** Comparing the optimum concentrations of the four extraction methods mentioned previously (0.4 or 0.2 mg/mL) revealed no significant difference in cell growth between any methods. $n$ = 6 distinct samples; statistical significance was calculated by one-way ANOVA and is indicated for $p > 0.2$ (n.s.).

*Supplementary Figure 2: Assessing additional potential albumin alternatives*

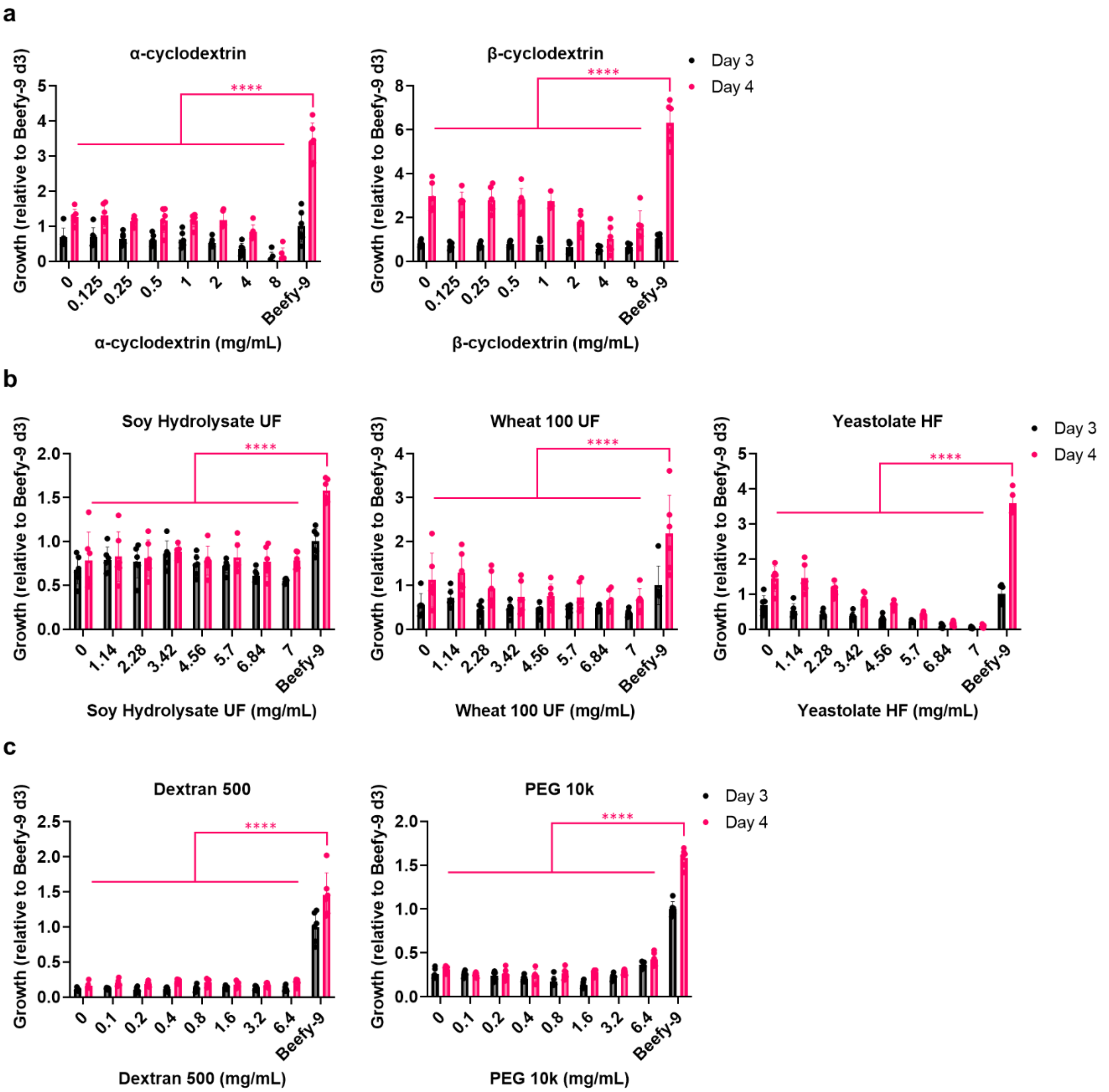


**Figure S2: Short-term growth of BSCs with other potential albumin alternatives**. **(a)** Cyclodextrins showed no improvement over cell growth compared with B8, and significantly reduced growth compared with Beefy-9. Analysis performed via Presto Blue metabolic assay on days 3 and 4 of growth. $n = 3$ distinct samples; statistical significance was calculated by one-way ANOVA for day 4 samples compared with Beefy-9 and is indicated for $p < 0.0001$ (****). **(b)** Supplier-provided plant protein hydrolysates showed no improvement over cell growth compared with B8, and significantly reduced growth compared with Beefy-9. Analysis performed via Presto Blue metabolic assay on days 3 and 4 of growth. $n = 3$ distinct samples; statistical significance was calculated by one-way ANOVA for day 4 samples compared with Beefy-9 and is indicated for $p < 0.0001$ (****). **(c)** Dextran 500 showed no improvement B8. PEG 10k showed minimal improvement at high concentrations. Both conditions showed significantly reduced growth compared with albumin-containing Beefy-9. Analysis performed via FluoReporter dsDNA quantification after days 3 and 4 of growth. $n = 6$ distinct samples; statistical significance was calculated by one-way ANOVA for day 4 samples compared with Beefy-9 and is indicated for $p < 0.0001$ (****).

*Supplementary Figure 3: Immunostaining of proliferative cells*

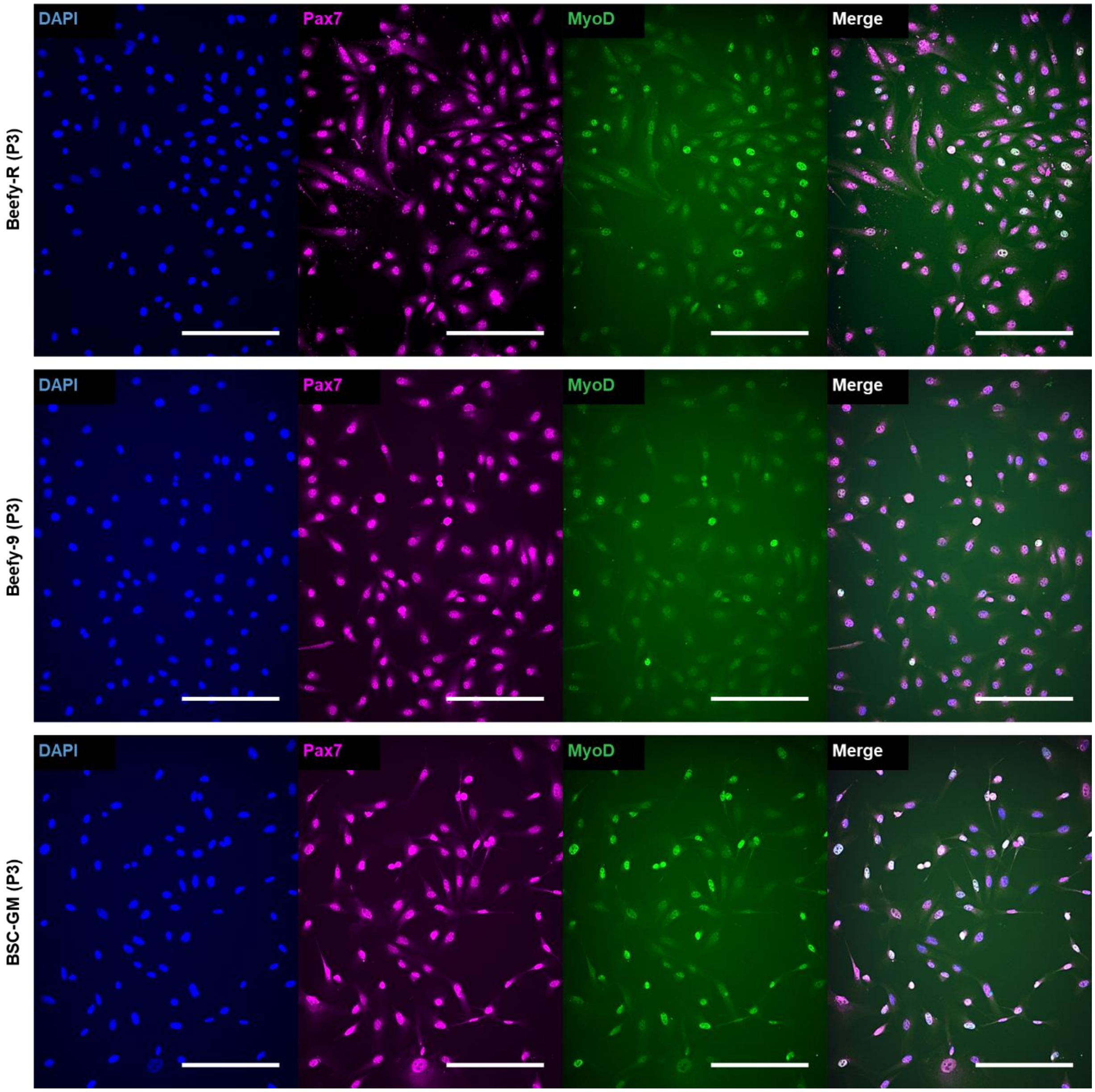


**Figure S3: Immunostaining of proliferative BSCs (p3) in various media.** Staining for nuclei (DAPI, blue), Pax7 (magenta), and MyoD (green). The results showed ubiquitous staining for Pax7 and heterogeneous staining for MyoD for all media types. Qualitative comparison showed a higher degree of bright MyoD staining for BSC-GM cells compared with Beefy-R and Beefy-9, which supports qPCR data seen in Fig. 4. Scale bars are 200 µm.

*Supplementary Figure 4: Example of Pax7 quantitation method*

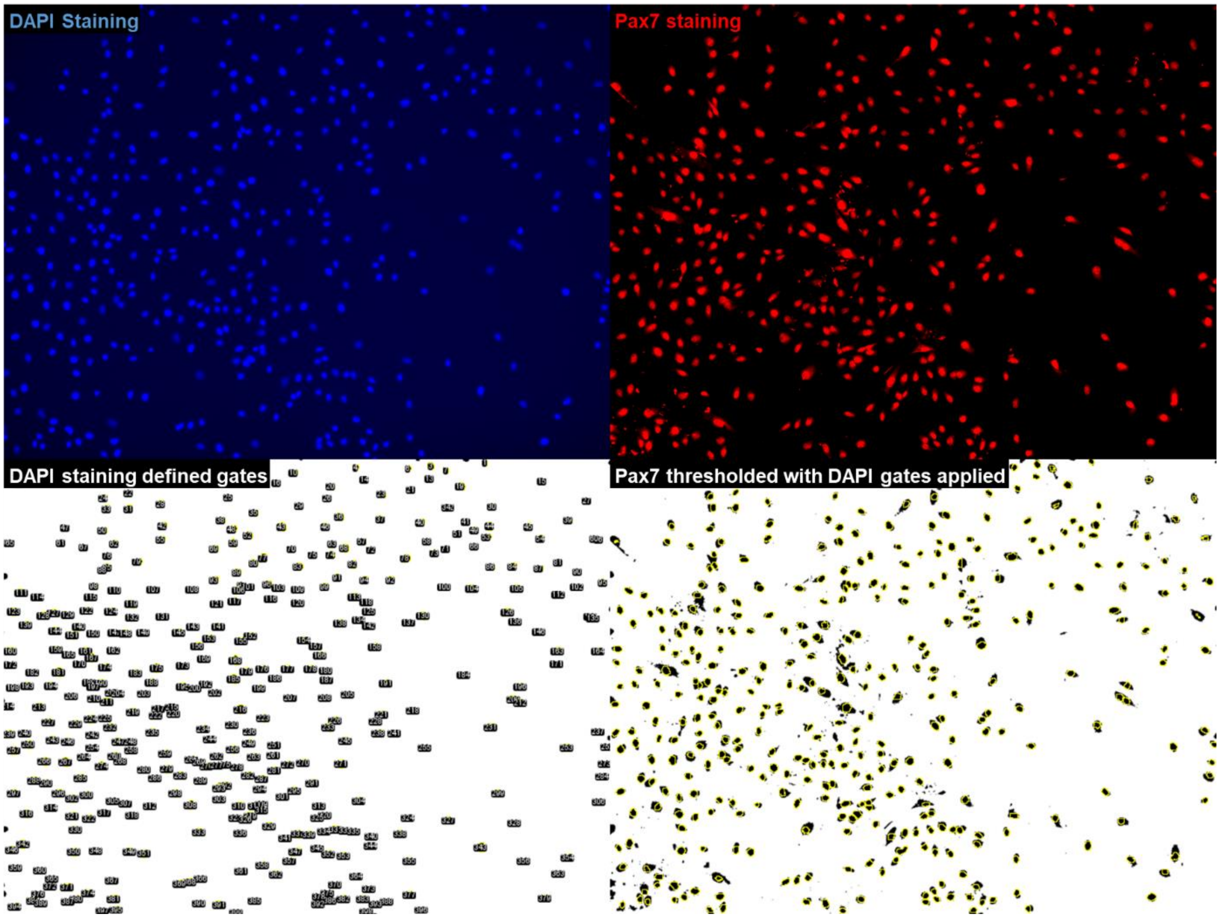


**Figure S4: Example Pax7 quantitation.** Example image represents one of 12 that was used for calculating Pax7-positive percentage in all media types. Top left: DAPI (nuclear) staining. Bottom left: DAPI staining converted to binary (black and white) and used to define nuclear regions. Top right: Pax7 staining. Bottom right: Pax7 staining with threshold applied and DAPI regions applied (yellow circles).

*Supplementary Figure 5: Immunostaining of differentiated cells*

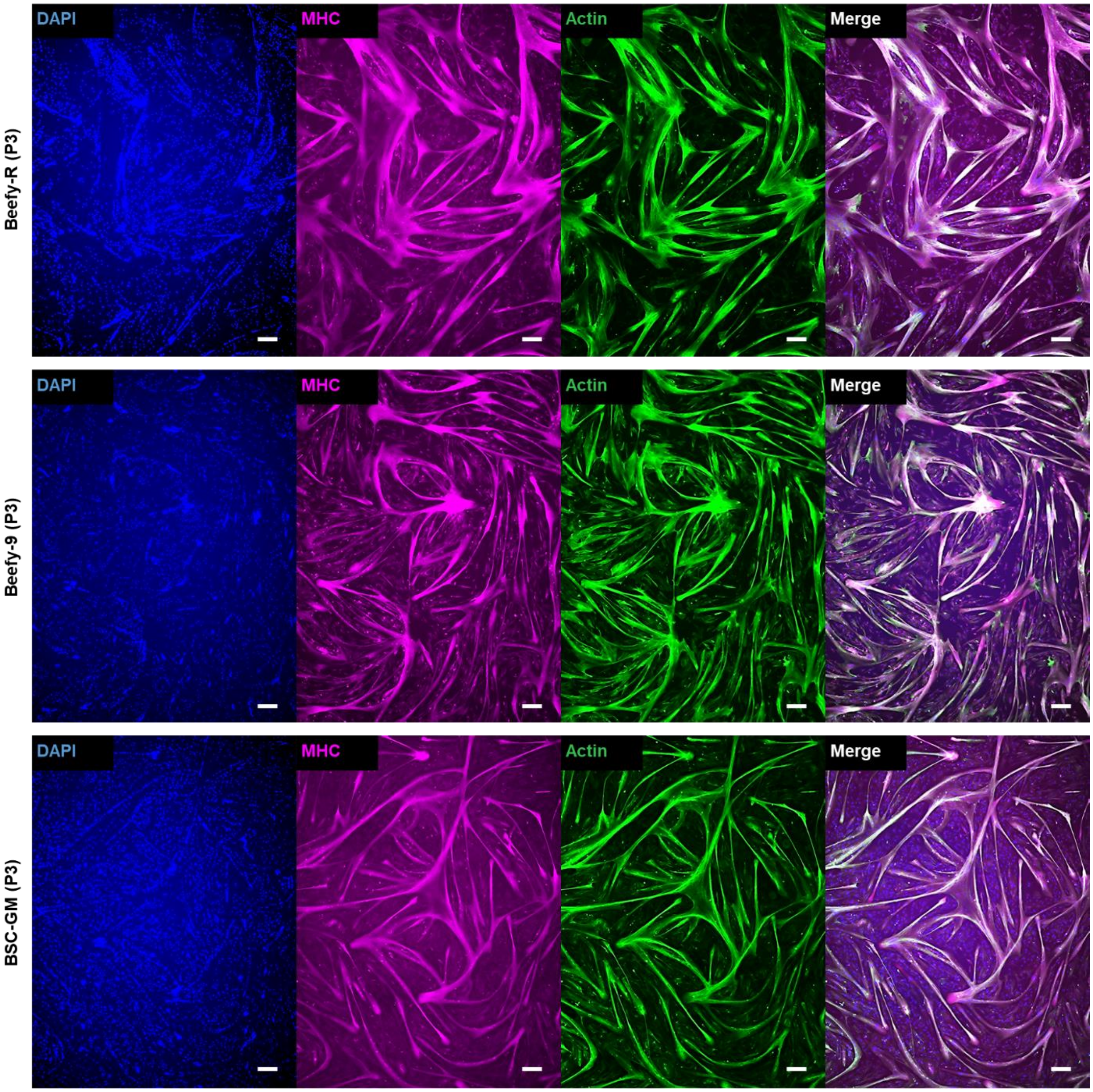


**Figure S3: Immunostaining of differentiated BSCs (p3) in various media.** Cells were differentiated for two days. Staining for nuclei (DAPI, blue), Myosin Heavy Chain (MHC; magenta), and Actin (green). Results showed robust myotube formation for all media types. Scale bars are 200 µm.

*Supplementary Figure 6: Example of fusion index calculations*

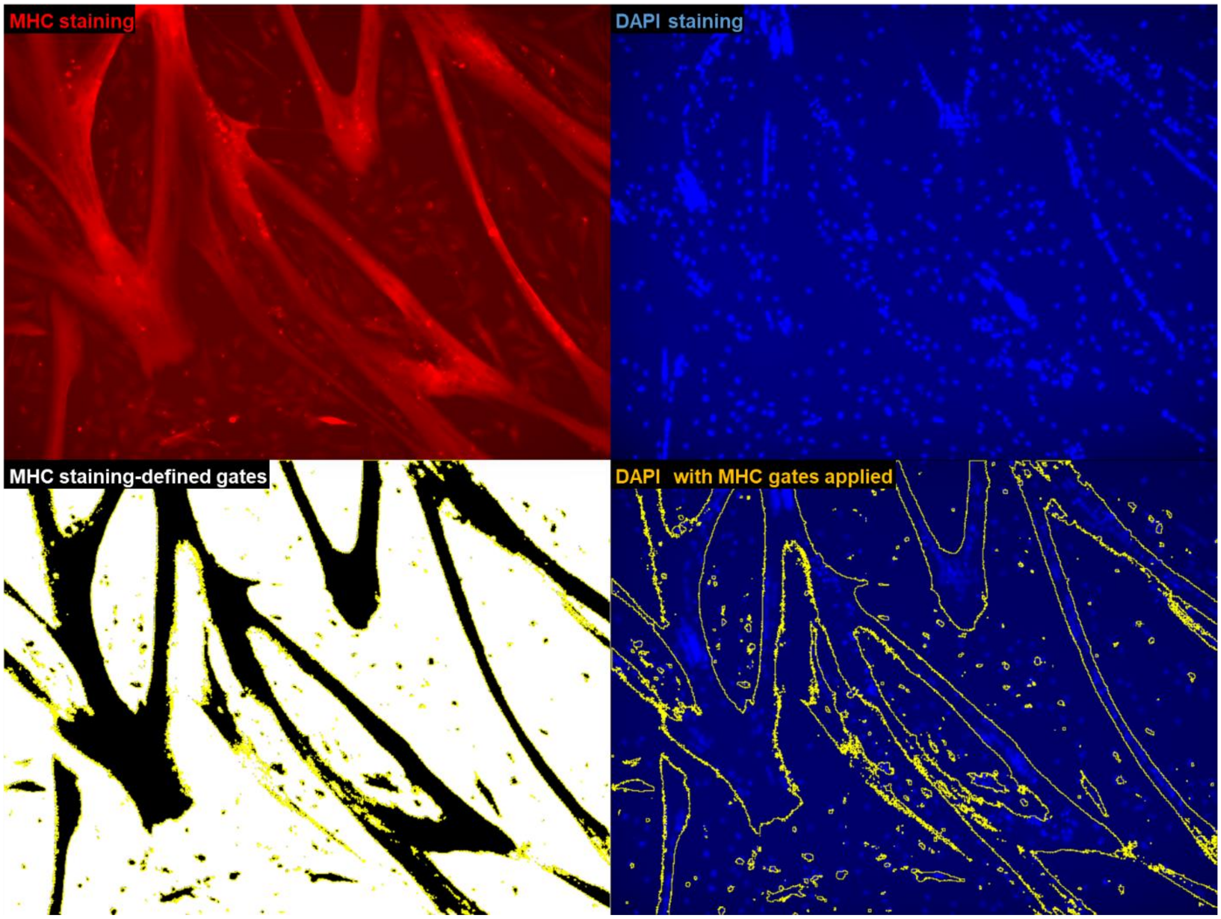


**Figure S4: Example fusion analysis.** Example image represents one of 12 that was used for calculating fusion index in all media types. Top left: MHC staining. Bottom left: MHC staining with threshold applied and used to define region of interest (yellow outline). Top right: DAPI (nuclear) staining. Bottom right: DAPI staining with region of interest applied. This image was used to calculate the ratio of nuclei within yellow-defined myotube regions compared with total nuclei.

*Supplementary Figure 7: Lipid droplet accumulation in various media*

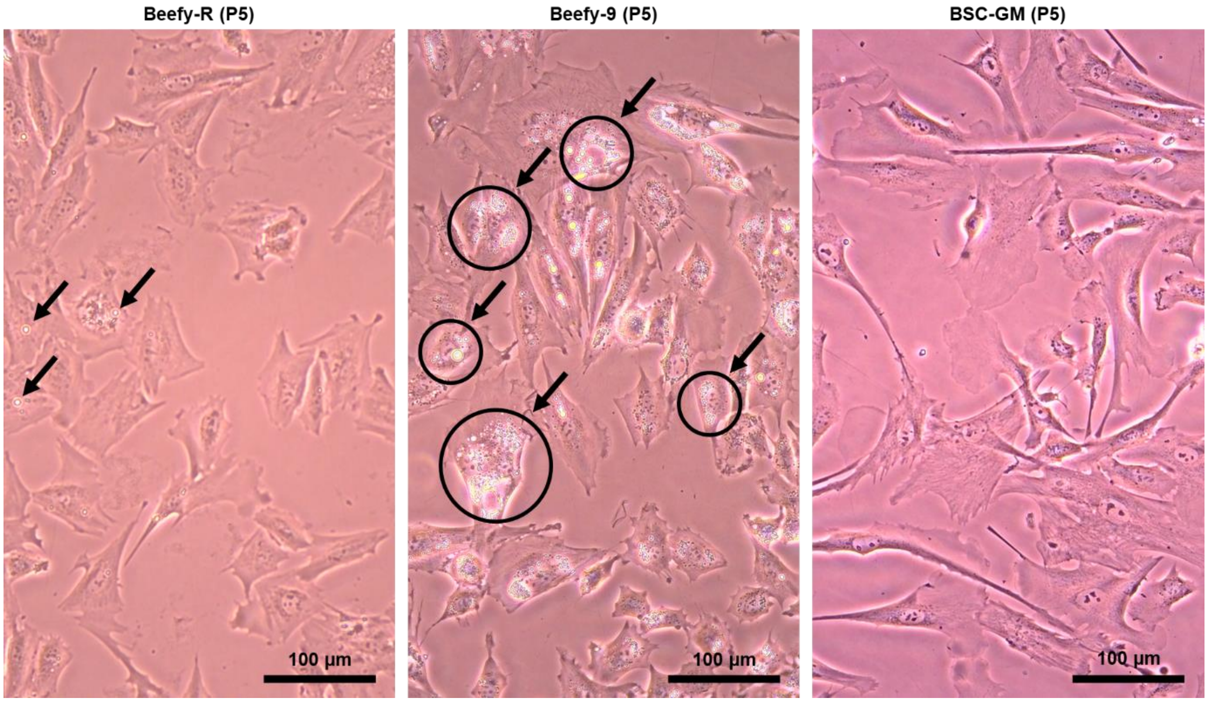


**Figure S7: Lipid accumulation in BSCs cultured in various media**. BSCs (P5) from various media conditions were imaged via brightfield microscopy to observe lipid droplet accumulation, which appears as bright circles within cells. Results showed that while some lipid droplet formation still occured in Beefy-R (examples indicated by black arrows; left image), it is substantially reduced compared with Beefy-9, which showed substantial lipid accumulation and the formation of large clusters of lipid droplets in cells (examples indicated by black arrows & black circles; middle image). BSC-GM controls showed very little to no aberrant lipid accumulation in cells. Scale bars are 100 µm.

*Supplementary Protocol 1: RPI and Beefy-R preparation*

**RPI Preparation:**

1. Grind rapeseed protein cakes (e.g., with a coffee grinder) to a finely ground meal.
2. Resuspend protein meal in DI water (1:10 w/v) and adjust pH to 12.5 with 5M NaOH.
3. Mix the protein meal slurry on a stir-plate for 1 hour and room temperature.
4. Centrifuge the protein meal slurry at 15,000 g for 10 minutes at room temperature.
5. Collect the supernatant and adjust pH to 4.5 with 6M HCl to precipitate proteins.
6. Centrifuge the protein mixture at 15,000 g for 10 minutes at room temperature.
7. Pour off the supernatant and keep the protein pellet.
8. Resuspend the protein pellet in the same volume of DI water that was used in the initial extraction and adjust pH to 12.5 with 5M NaOH.
9. Mix on a stir-plate until protein pellet is fully dissociated. Note: it may be challenging to dissociate the pellet. If so, a combination of stir-plate mixing, and mechanical dissociation can help (e.g., with a plastic or metal weighing spatula).
10. Adjust pH to 7.2 with 6M HCl.
11. Centrifuge the protein solution at 15,000 g for 30 minutes at room temperature.
12. Filter supernatant through a 10 µm filter.
13. Centrifuge at 45,000 g for 3 hours at 4°C.
14. Filter through 0.22 µm sterile filter.
15. Concentrate protein solution through a 3kDa MWCO filter. Note: concentrating the solution 50-100x will typically result in a concentration at least 50 mg/mL, which will enable dilution to the working concentration of 50 mg/mL. In this study, concentration was performed via ultrafiltration through a 3 kDa cutoff membrane via centrifugation at 4,500 g overnight at 4°C; however, other concentration methods can be used.
16. Sterilize the concentrated RPI through an 0.45 µm$^2$ filter.
17. Collect the concentrated protein solution and quantify (e.g., via BCA or Bradford assay).
18. Dilute RPI to 50 mg/mL with sterile DI water.
19. Aliquot RPI, snap-freeze with liquid nitrogen, and store long-term in -80°C.

**Beefy-R Preparation:**

1. Prepare B8 in one of two ways:
    a. Add HiDef-B8 aliquots from Defined Bioscience to DMEM/F12 according to the manufacturer's instructions.
    b. Prepare B8 according to instructions given in the supplementary information of the medium's original publication (Kuo et al., 2020).
2. Thaw RPI at 4°C, on ice, or under cool running tap water.
3. Prepare a tube with the necessary volume of B8 for scheduled cell feeding.
4. Add RPI at a concentration of 0.4 mg/mL.
5. Feed cells using standard procedures.

**Beefy-R final composition:**

<table>
<tr><th>Component</th><th>Per Liter</th><th>Supplier (In-house B8)</th><th>Supplier (HiDef-B8)</th></tr>
<tr><td>DMEM/F12</td><td>1 L</td><td colspan="2">ThermoFisher #11320033</td></tr>
<tr><td>L-ascorbic acid 2-phosphate</td><td>200 mg</td><td>Sigma #49752-10G</td><td rowspan="7">Defined Bioscience #LSS-201</td></tr>
<tr><td>Insulin</td><td>20 mg</td><td>Sigma #91077C-100MG</td></tr>
<tr><td>Transferrin</td><td>20 mg</td><td>Sigma #T3705-1G</td></tr>
<tr><td>Sodium selenite</td><td>20 µg</td><td>Sigma #S5261-10G</td></tr>
<tr><td>Fibroblast growth factor (FGF-2 or FGF2-G3)</td><td>40 µg</td><td>PeproTech #100-18B</td></tr>
<tr><td>Neuregulin (NRG1)</td><td>100 ng</td><td>PeproTech #100-03</td></tr>
<tr><td>Transforming growth factor beta-3 (TGFβ-3)</td><td>100 ng</td><td>R&D Systems #8420-B3-005/CF</td></tr>
<tr><td>Rapeseed protein isolate (RPI)</td><td>400 mg</td><td colspan="2">Prepared in-house</td></tr>
<tr><td>Antibiotic-antimycotic (100x)</td><td>10 mL</td><td colspan="2">ThermoFisher #1540062</td></tr>
</table>
